# DNA Gap Repair-Mediated Site-Directed Mutagenesis is Different from Mandecki and Recombineering Approaches

**DOI:** 10.1101/313155

**Authors:** George T. Lyozin, Luca Brunelli

## Abstract

Site-directed mutagenesis allows the generation of mutant DNA sequences for downstream functional analysis of genetic variants involved in human health and disease. Understanding the mechanisms of different mutagenesis methods can help select the best approach for specific needs. We compared three different approaches for *in vivo* site-directed DNA mutagenesis that utilize a mutant single-stranded DNA oligonucleotide (ssODN) to target a wild type DNA sequence in the host *Escherichia coli* (*E. coli*). The first method, Mandecki, uses restriction nucleases to introduce a double stranded break (DSB) into a DNA sequence which needs to be denatured prior to co-transformation. The second method, recombineering (recombination-mediated genetic engineering), requires lambda *red* gene products and a mutant ssODN with homology arms of at least 20 nucleotides. In a third method described here for the first time, DNA gap repair, a mutant ssODN targets a DNA sequence containing a gap introduced by PCR. Unlike recombineering, both DNA gap repair and Mandecki can utilize homology arms as short as 10 nucleotides. DNA gap repair requires neither *red* gene products as recombineering nor DNA denaturation or nucleases as Mandecki, and unlike other methods is background-free. We conclude that Mandecki, recombineering, and DNA gap repair have at least partly different mechanisms, and that DNA gap repair provides a new, straightforward approach for effective site-directed mutagenesis.

## Introduction

Site-directed mutagenesis can be achieved by replacing a wild type DNA sequence with a premade mutant DNA sequence, i.e., the targeting vector. Mutant single-stranded DNA oligonucleotides (ssODNs) are the most useful targeting vectors because they are commercially available and relatively inexpensive. For DNA-DNA recognition, two regions at the ends of the targeting DNA (homology arms) need to be homological to the flanks of the site to be mutated in wild type DNA (targeted DNA). The region of the targeting vector between the homology arms contains the mutation(s) to be introduced into targeted DNA. The approach is called DNA integration when the targeted DNA sequence is intact (Fig. 1). Due to significant background cell contamination, a mutant isolation procedure is usually required in this approach. In DNA repair, targeted DNA is discontinuous due to the introduction by site specific endonucleases of at least one DSB between the homology arms. A gap is produced when a segment of the targeted DNA is deleted by two or more DSBs (Fig. 1). In DNA integration, DNA-DNA recognition relies only on homology arms, but in DNA repair there is also the need for site recognition by nucleases. Thus, one might expect DNA integration to be not only easier to implement but also more specific. Recombineering is a technique to manipulate DNA in *E. coli* that relies on short homology sequences and requires bacteriophage lambda *red* gene products for its functionality. This applies to both DNA integration and DNA repair^1^. Recombineering is particularly well suited for single-copy, high capacity plasmids such as bacterial artificial chromosomes (BACs). However, with high copy number plasmids only one copy is typically modified, and segregation of mutants is challenging. In addition, induction of *red* genes can lead to the formation of high molecular weight plasmids^2^.

**Fig. 1.**
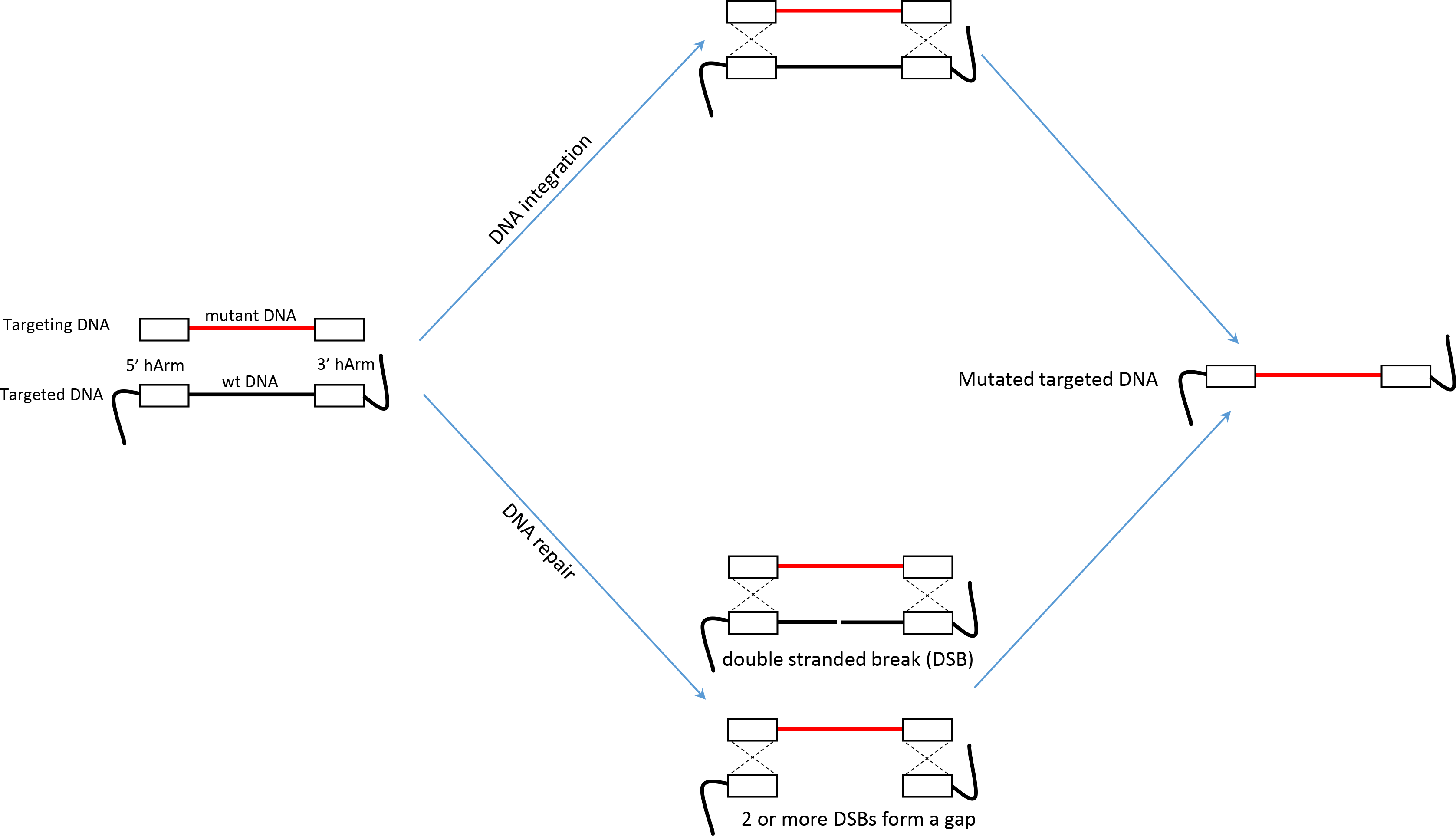
Different ways of engineering DNA. With DNA integration, targeted DNA remains intact between the homology arms (hArms). With DNA (gap) repair, there is one or more DSBs, forming a gap in the DNA sequence between the homology arms.

The Mandecki method utilizes denatured plasmid DNA linearized with restriction endonucleases and a ssODN targeting vector (Fig. 2a)^3^, but does not require phage lambda transgenes. There is no problem with mutant segregation because the *E. coli* host does not contain wild type plasmid, and mutants are generated in cells transformed with linearized plasmid and mutant ssODN. Thus, this approach seems better suited to mutate high-copy number plasmids.

**Fig. 2.**
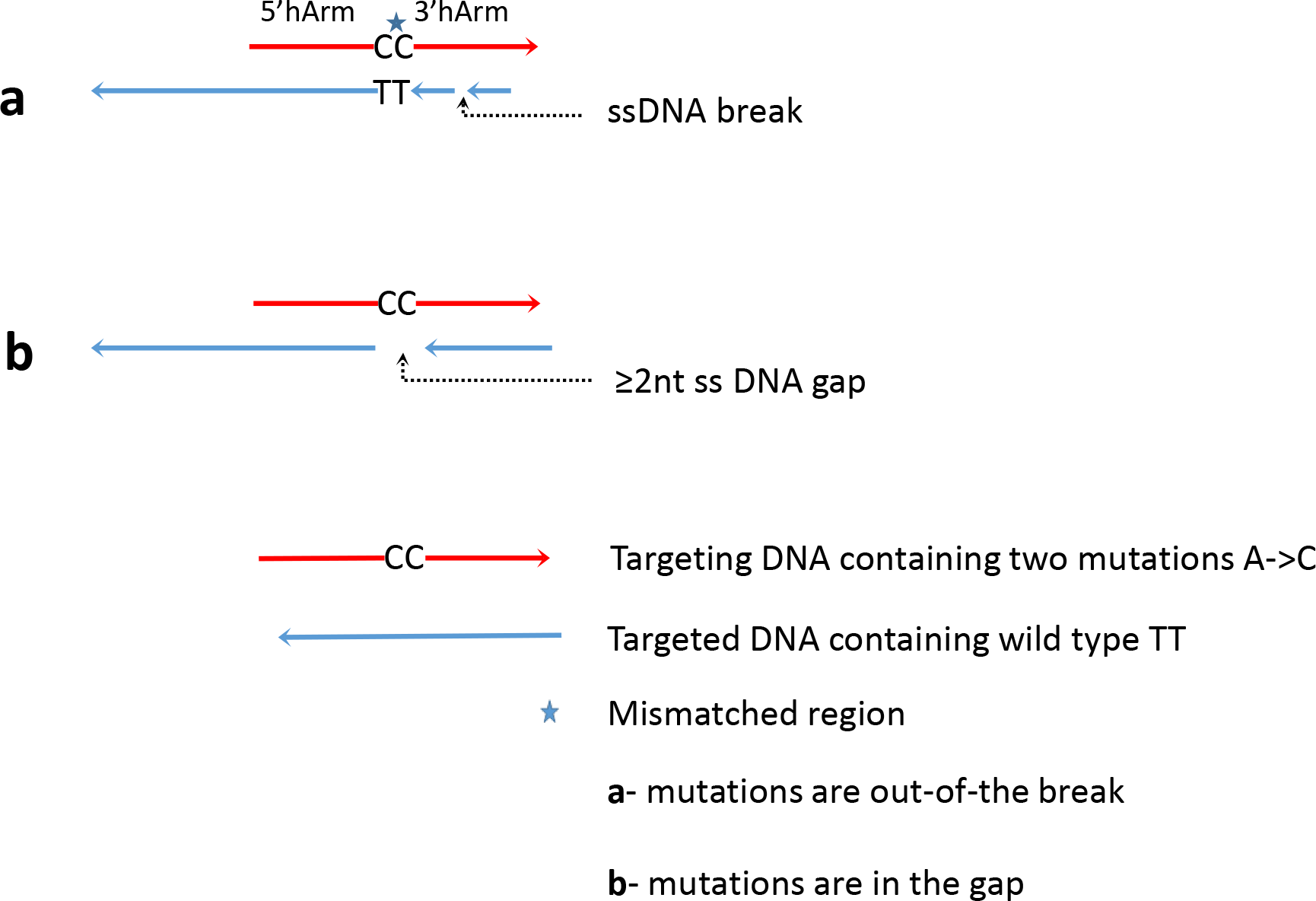
Two possible localizations of a mutation in DNA repair. A. The mutation is not located in the DSB(s), but it is in the homology arms where it forms a mismatched heteroduplex with the targeted DNA. B. The mutation is located within the gap. The homology arms of the targeting DNA perfectly match the targeted DNA. Only the strand of targeted DNA complementary to ssODN is shown.

Here we studied a ‘pure’ DNA gap repair approach in *E. coli* in which the sequence to be mutated is always pre-deleted from wild type targeted DNA. Upon DNA gap repair, the targeting and targeted DNA form perfectly matched hetero-duplexes (Fig. 2b). To mutate a certain number of nucleotides, at least the same number of nucleotides need to be deleted from targeted DNA. With relatively short plasmids (<8 kb), this can be conveniently achieved by using the polymerase chain reaction (PCR) to amplify only the DNA segment that is not undergoing mutagenesis, thus generating a linear DNA construct with a gap (Fig. 3). Some important features of recombineering and Mandecki that will be compared with DNA gap repair are summarized in Table 1.

**Table 1.**
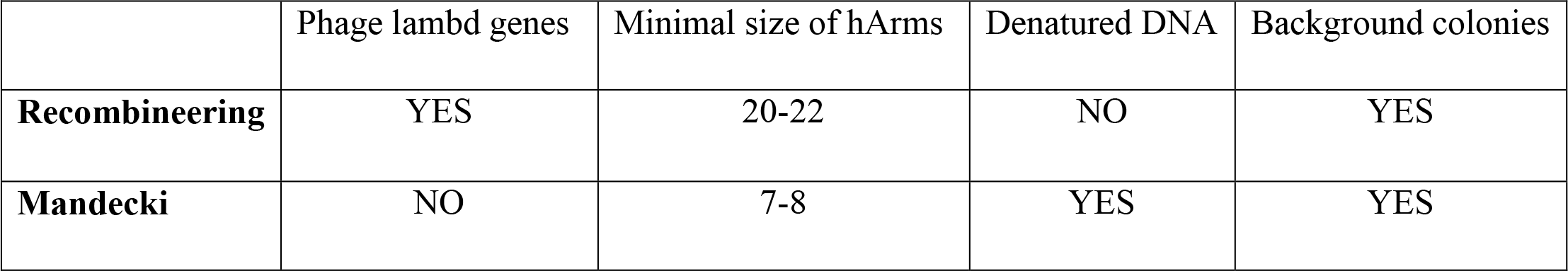
Features of recombineering and Mandecki approaches

|  | Phage lambd genes | Minimal size of hArms | Denatured DNA | Background colonies |
| --- | --- | --- | --- | --- |
| **Recombineering** | YES | 20-22 | NO | YES |
| **Mandecki** | NO | 7-8 | YES | YES |

**Fig. 3.**
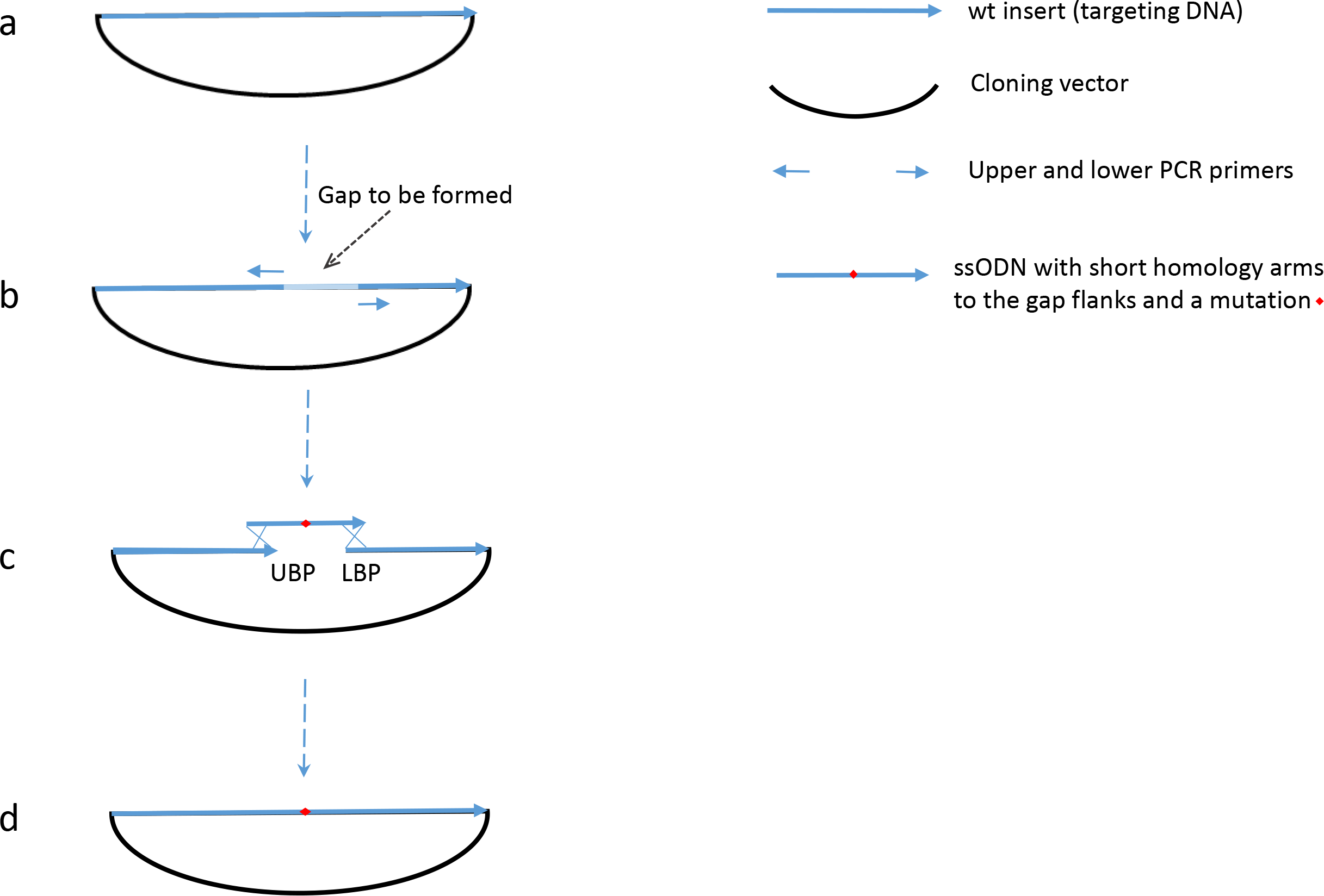
Site-directed mutagenesis mediated by DNA repair in a gap introduced by PCR. A. Original DNA construct containing wild type insert cloned in a vector. B. Amplification by PCR of the DNA construct which “deletes” the sequence between the 5’ primer ends of the insert. C. The gap formed between the upper break point (UBP) and lower break point (LBP) is repaired by a mutant ssODN. D. DNA repair of the gap with the mutant ssODN leads to the insertion of the mutation in the construct.

## Materials and Methods

The approximately 8,500 base pair (bp) wild type construct contained 714 bp wild type genomic sequence of the mouse *Tnnt2* gene cloned in our compact P1 phage-based vector^4^. The insert was obtained by PCR using primers fPhoTnT2BAC_ 30679 and rPHOTnT2BAC_31392 (Table 2) and template RP23-389C7 BAC DNA. The insert sequence corresponds chromosome 1, assembly GRCm38.p4 with the coordinates: 135847355-135848068 (NCBI Reference Sequence: NC_000067.6). The insert was blunt-end ligated to an approximately 7,800 bp PCR product of the vector DNA generated with primers f_-TkRV_8976 and r_-TkRV_7573 (Table 2). Targeted DNA was obtained from the wild type construct described above and primers fMAMCon8070PAG and r91TnTCon7950PAG (Table 2). The 21-nucleotide gap TGCCACCCAAGATCCCCGATG representing a part of the wild type construct, which was not amplified, was formed by the 5’-end boundaries of the primers. All targeting ssODNs with the sequences showed in Table 2 inserted the first nucleotide of the gap, i.e., T (showed in capital case). All ssODNs were ordered from Thermo/Life Sciences and were desalted except fMAMCon8070PAG, r91TnTCon7950PAG and 2×30+1 which were PAAG purified.

**Table 2.**
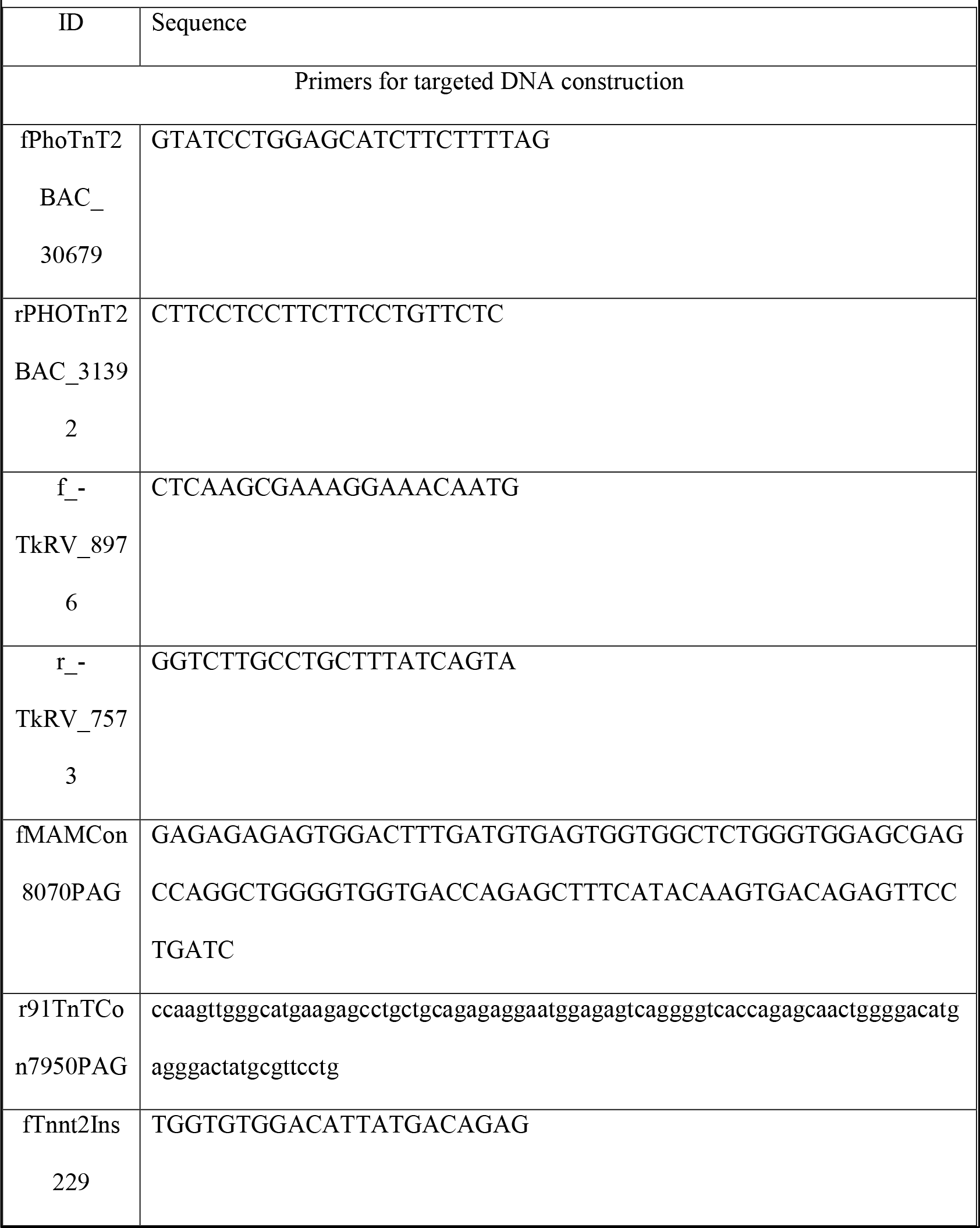
PCR primers and targeting ssODNs used in experiments

### Chemicals and enzymes

Chemicals were purchased from BioBasic (Markham, Ontario, Canada) or ThermoFisher (Waltham, MA). PrimeSTAR GXL polymerase was purchased from Takara Bio USA (Mountain View, CA).

### Plasmids

Wild-type *Tnnt2* BAC, 209.7 kb RP23-389C7 BAC clone (http://bacpac.chori.org/).

### DNA isolation and purification

Plasmid DNA was extracted as previously described^5^, but the composition of the non-ionic detergent isolation buffer was changed for large scale plasmid DNA extraction as follows: 1.75 M NH_4_Cl, 50 mM EDTA, and 0.15% IGEPAL CA-630, RNase A, and lysozyme. Some DNA sequencing samples were PCR products prepared with a new express method (Lyozin and Brunelli, unpublished observation).

### DNA sequencing

DNA sequencing was performed at the University of Nebraska Medical Center DNA Sequencing Core Facility using the ABI BigDye Terminator Cycle Sequencing Kit v3.1 according to the manufacturer’s instructions on an Applied Biosystems 9800 Fast Thermal Cycler. The sequence fragments were detected on an ABI 3730 DNA Analyzer. Samples were then analyzed and base-called by Applied Biosystems DNA Sequencing Analysis Software V5.2 (Applied Biosystems, Foster City, CA).

### Bacterial transformation

Electro-competent cells were prepared as described previously^6^. DNA was electroporated at 17,000-18,000 V/cm at time constant 7.5 msec using BioRad Gene Pulser Xcell into 10 μl competent cells. Native targeted DNA before electroporation was concentrated and desalted using Amicon Ultracel-100K filter units (Millipore Sigma, Burlington, MA). 2 μl electroporated DNA contained 50 ng of the native targeting DNA and 10 pms ssODN. Denatured DNA was prepared as described by Mandecki, and concentrated and desalted as described above. 2-3 μl DNA electroporated contained 50 ng of thermally denatured DNA and additional 20 pms ssODN added after ultrafiltration.

Alkali to denature DNA was in a solution of 0.2 M NaOH, 5 mM EDTA, 100 mM NaCl. After denaturing DNA was precipitated with EtOH and washed with 70% EtOH, the DNA pellet was dissolved in electroporation buffer^6^. The alkali denatured DNA concentration was measured with DeNovix which clearly detected the hyperchromic shift due to DNA denaturation. The denatured DNA was stored at 4 °C and used for transformation without delay. CaCl_2_ transformation of JM83 cells obtained from Addgene with thermally denatured targeted DNA and ssODN was performed as described by Mandecki except for using SOC media instead of 2x YT for Amp^r^ marker expression. Mix&Go transformation reagents and Mix&Go JM108 chemically competent cells were obtained from Zymo Research (Irvine, CA) and used according to the manufacturer’s instructions. Sterile 10 mM MgCl_2_ solution was used for bacterial culture dilution.

### Recombineering

Mini-λ provided by Dr. Shyam K. Sharan (NCI-Frederick) was introduced in DH10B T1 resistant cells obtained from New England Biolabs (Ipswich, MA) as described previously^7^.

### PCR

PCR mix to generate the targeted DNA with a gap contained 0.1 μM primers; 0.2 mM dNTP; 5% DMSO, 2.5% glycerol, 0.5 ng/ml vector template DNA and 10 U PrimeSTAR GXL polymerase. Because the primers are long, PCR included 35 two-step cycles at 96°C for 15” and at 70°C for 9’30”. The PCR products were treated with DpnI as previously described^6^, then precipitated with isoPrOH and washed with 70% EtOH. The PCR mix is very efficient and specific and no further purification for DNA gap repair experiments was required.

The PCR mix used to generate the targeted DNA with terminal direct repeats (TDR) contained 0.02-0.033 μM overlapping primers (Fig. 6, legend); 0.2 mM dNTP; 0.5 M betaine; 50-300 pg/μl template DNA and 12.5 U/ml PrimeSTAR GXL polymerase. PCR included 20-25 cycles (96°C for 15”, 55°C for 10” and 65°C for 4’30”). Specific PCR product yield was often low. Increasing DNA template concentration, or decreasing primer concentration, primer annealing time and number of cycles helped obtaining specific PCR products. However, even when the majority of PCR mixes contained the expected products, gel electrophoresis purification was required for *in vivo* experiments.

## Results and Discussion

### DNA gap repair requires highly efficient DNA intracellular delivery

In most recombineering applications, targeted DNA resides in the *E. coli* host, and thus only targeting DNA needs to be delivered into *E. coli* for recombination. However, in the Mandecki approach both targeting and targeted DNA need to be delivered into the *E. coli* host. This method is therefore expected to be less efficient, and requiring cells highly competent for DNA uptake. In agreement with this, we were unable to detect any colonies with the co-electroporation of targeted DNA and ssODN with an efficiency of <10^8^ colony forming units (CFUs) per μg pUC18 plasmid DNA. Details on efficiency are provided in Fig. 4, legend.

**Fig. 4.**
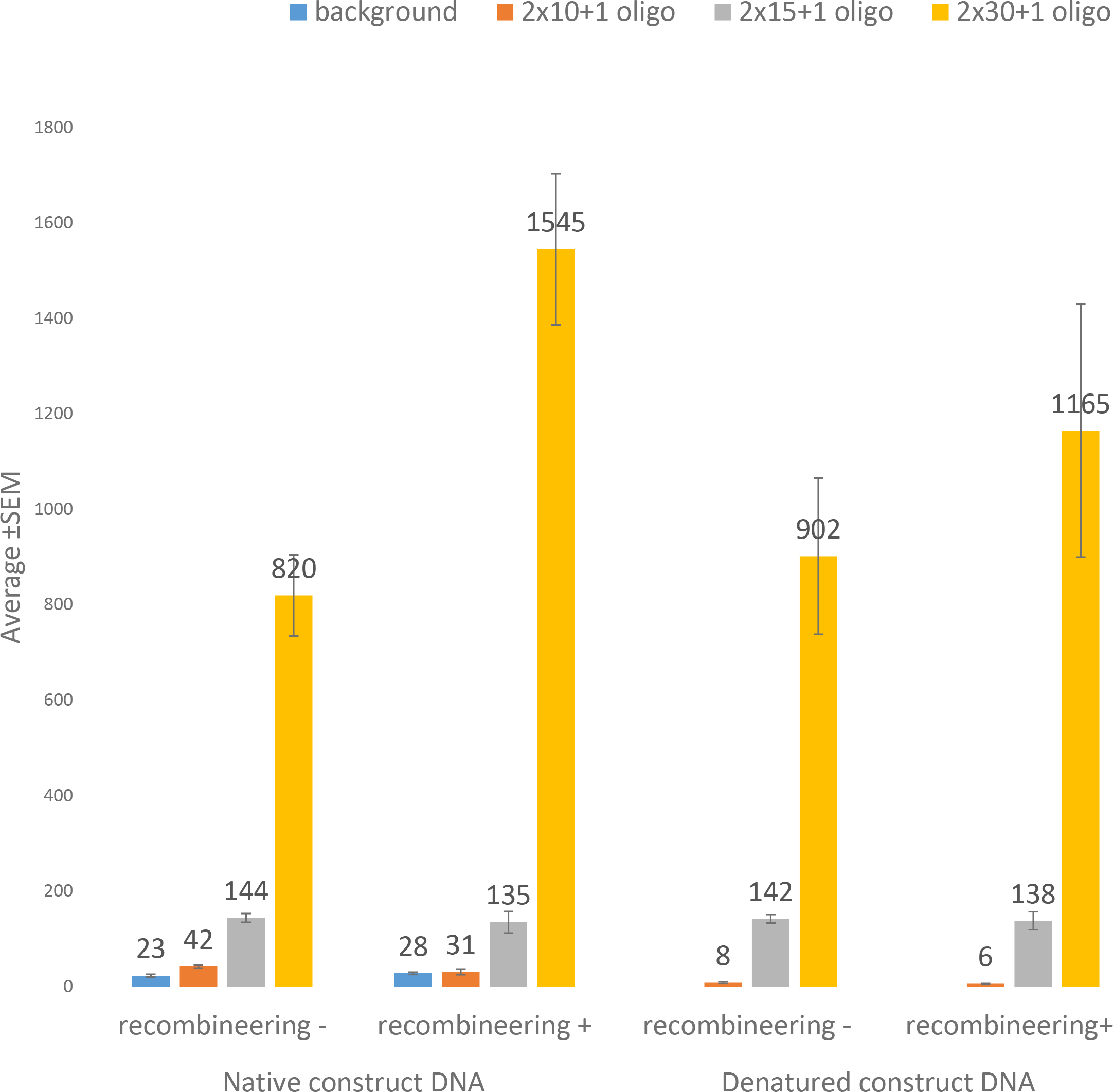
Colony numbers with DNA gap repair. DH10B cells with repressed (recombineering-) and de-repressed (recombineering+) lambda *red* genes were co-electroporated with 50 ng of either native targeted DNA (native construct DNA) or thermally denatured DNA (denatured construct DNA) and 10 pms ssODNs with different homology arms: 10 nt (2×10+1); 15 nt (2×15+1); and 30 nt (2×30+1). All ssODNs were designed to insert T in the 21 bp gap of targeted DNA. The columns with error bars represent average colony numbers per plating. Background colony numbers were determined through the electroporation of native construct DNA. Electroporation efficiency of the cells were determined from triplicates and were equal to 3.3×10^9^ for repressed cells (recombineering-) and 2.2×10^9^ for de-repressed cells (recombineering+).

### DNA gap repair is driven by endogenous pathways independent of *red* genes

DNA integration requires recombineering genes products, while the minimal size of the homology arms is probably ~22 nucleotides^8, 9^. To determine whether recombineering genes products are required for DNA gap repair, we co-electroporated lambda phage lysogens with a targeting ssODN with homology arm length of either 10, 15 or 30 nucleotides to insert, as in the Mandecki report^3^, one nucleotide in the gap (indicated in Fig. 4 and 5 as 2×10+1, 2×15+1, and 2×30+1, respectively) (Table 2). We then counted the number of colonies with or without de-repression of lambda genes (leading to rapid intracellular accumulation of recombineering gene products). As shown in Fig. 4, increasing homology arm length led to higher colony numbers in both repressed and de-repressed conditions. Importantly, with 10 and 15 nucleotide homology arms, colony counts in both conditions were similar. With 30 nucleotide homology arms, the expression of recombineering genes led to doubling of the number of colonies. These results indicate that at least one of the three *red* genes promotes DNA gap repair when homology arm length is >15-30 nucleotides. Importantly, the expression of recombineering genes does not affect DNA gap repair using homology arms of 10 to 15 nucleotides.

**Fig. 5.**
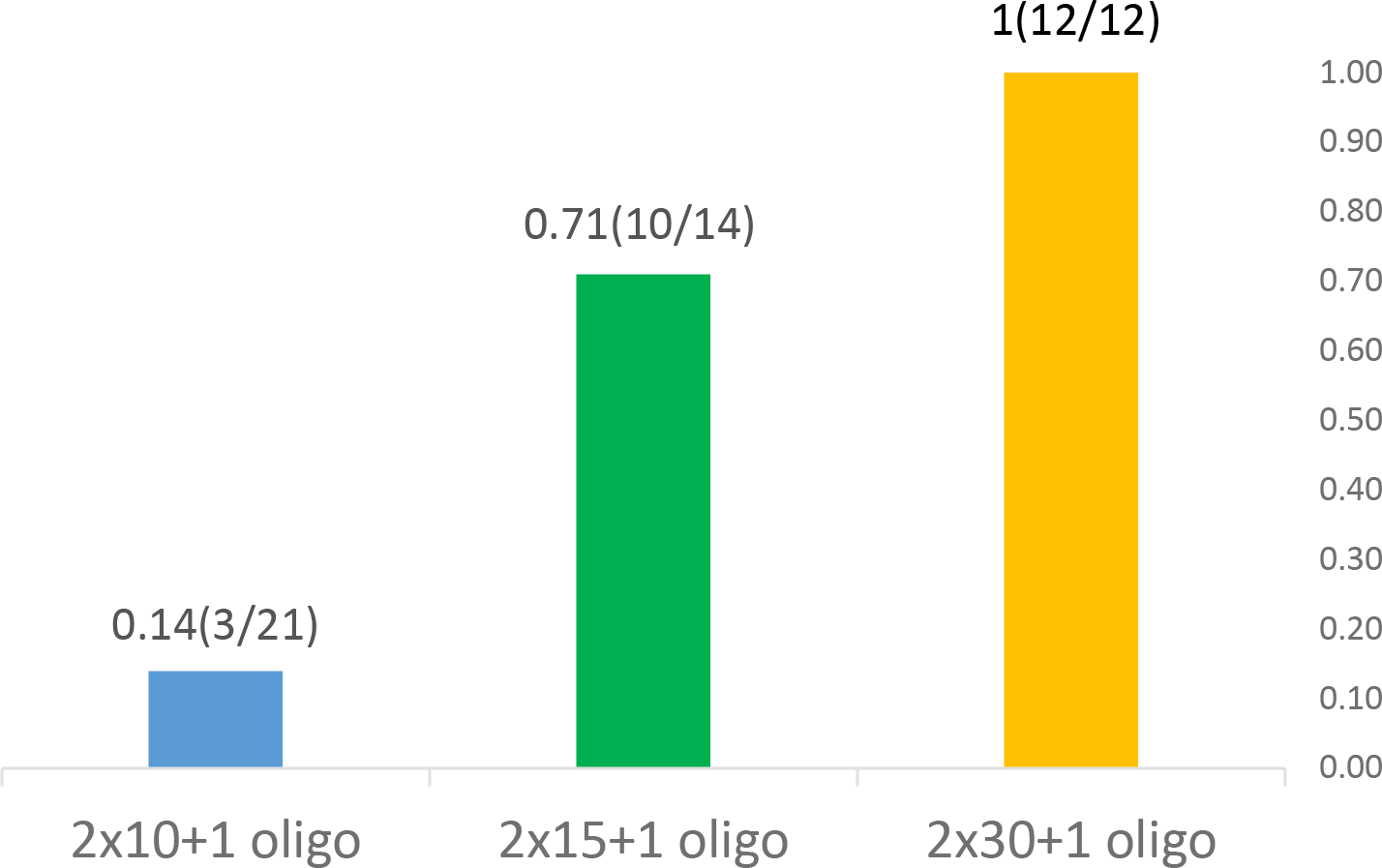
Relationship between homology arm length and frequency of mutant colonies. Three targeting ssODNs with different homology arms were used: 10 nt (2×10+1); 15 nt (2×15=1); and 30 nt (2×30+1). A number of colonies indicated in the denominator were sequenced, and the numbers of mutants confirmed by DNA sequencing represent the numerator of the fraction.

**Fig. 6.**
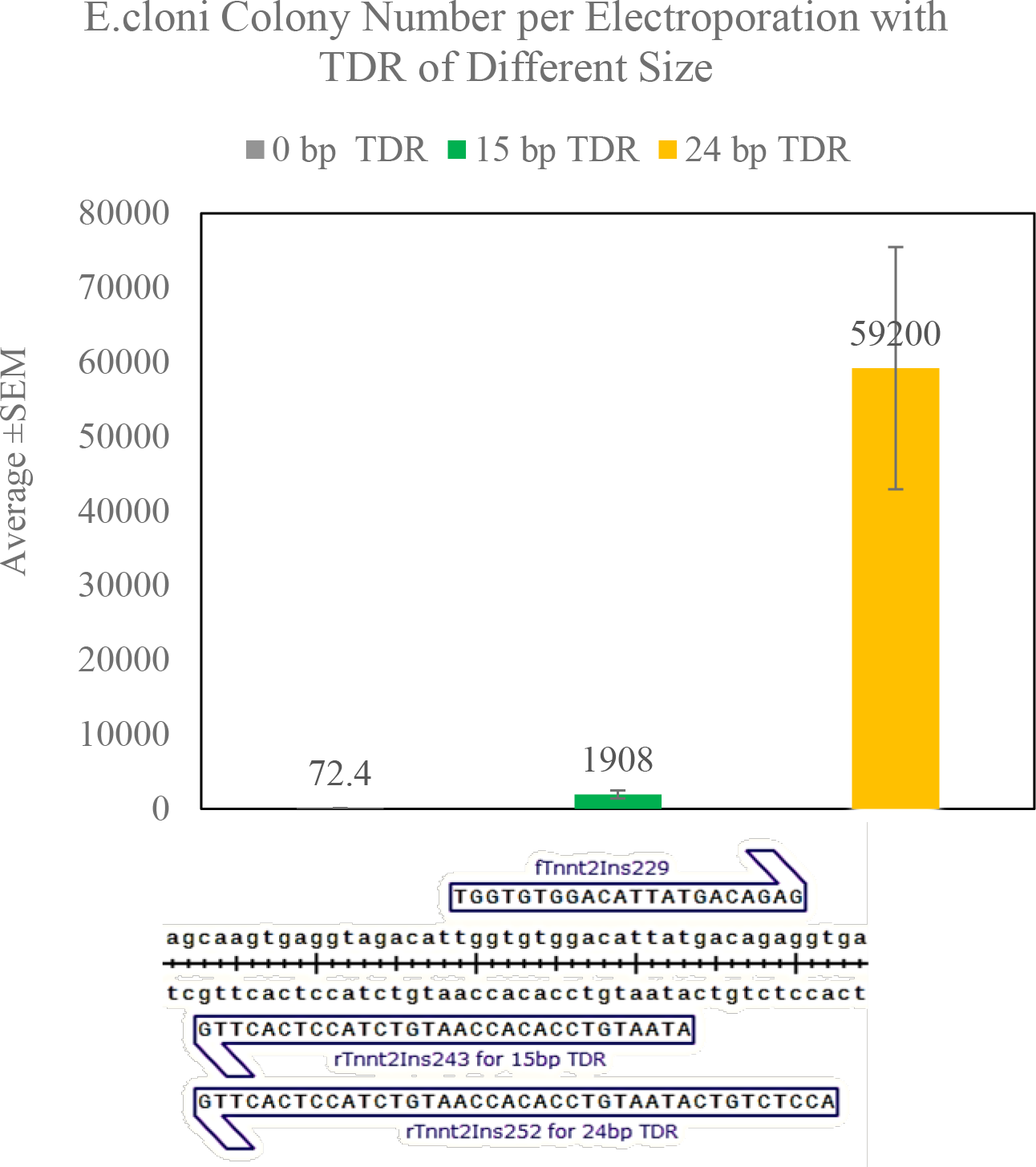
Relationship between the size of terminal direct repeats (TDR) and the frequency of background cells. Electrocompetent *E.cloni* 10G cells were purchased from Lucigen. 5 pg standard pUC plasmid DNA and 20 ng DNA with no TDR; 15 bp TDR; and 24bp TDR treated with DpnI and purified with gel electrophoresis, were electroporated into cells (3-5 replicas each). Electroporated cells were plated on Amp agar and the number of colonies was calculated. Efficiency of electroporation was 6×10^10^ and the total number of cells surviving electroporation was about 10^8^. Average number of colonies per electroporation and standard error of the mean are presented in the chart. Different size TDR DNA was generated using overlapping forward and reverse primers as shown below the figure.

### DNA denaturation is not required for DNA gap repair

Mandecki’s buffer for DNA thermal denaturation contains salts, precluding the use of denatured DNA solutions in electroporation because of their high conductivity. We used membrane ultrafiltration for desalting DNA solutions prior to electroporation. Due to partial renaturation, the structure of denatured DNA could change during this about 15-minute procedure compared to denatured DNA stored at 4 °C until transformation as in the Mandecki method. However, partial DNA renaturation could not be excluded in the Mandecki method because DNA transformation involves a three-minute incubation at 37 °C in the presence of 50-100 mM CaCl_2_. Although structural DNA changes in these types of experiments can proceed to varying degrees, we tested thermally denatured DNA in our experiments because some fraction of particular recombination-active structures could persist after thermal denaturation and DNA desalting. As shown in Fig. 4, DNA denaturation significantly decreased colony counts in both recombineering-induced and non-induced cells when the gap was repaired with 10-nucleotide homology arm ssODN, but there was no difference with 20 and 30-nucleotide homology arm ssODNs. Denatured targeted DNA might be less active in DNA gap repair and increasing homology arm length might restore activity by promoting DNA renaturation. In agreement with this, co-transformation of commercial chemically-competent cells (DNA desalting is not required) with 40-nucleotide homology arm ssODNs and denatured DNA followed by ultrafiltration produced twice more colonies compared to experiments done without ultrafiltration. In similar experiments, no colonies were produced when DNA was denatured with alkali (see Materials and Methods). Alkali-denatured DNA did not produce any colony in electroporation experiments as well (data not shown). This evidence supports the conclusion that DNA denatured according to the Mandecki method changed its structure during ultrafiltration, and denatured DNA is not active in DNA gap repair.

### Efficiencies of Mandecki method and DNA gap repair

To correctly compare the efficiencies of different genetic engineering pathways in *vivo*, the number of recombinants should be normalized to the number of cells penetrated by recombining DNA (both targeting and targeted). However, these numbers are usually not available because even transformation efficiencies and fraction of competent cells are known only for covalently closed double stranded plasmid DNA^10^. Currently, efficiencies are calculated as the fraction of mutant to the total number of cells, frequently ignoring important differences in DNA uptake efficiency by cells. Mandecki used “efficiency of mutagenesis”, that is the fraction of mutant colonies among the total number of colonies (mutants plus non-mutants) on agar plates. However, this value does not provide information on how easily mutant colonies can be generated from the total number of original cells (this is the true efficiency). This fraction varied between 0.2% (2 mutants per 820 transformants) and 98% (1,330 mutants per 1,360 transformants). The latter was obtained with a 30 nucleotide homology arms ssODN inserting a nucleotide into thermally denatured DNA.

To compare DNA gap repair and Mandecki method efficiencies, we determined that cells surviving electroshock using a 30-nucleotide homology arms ssODN were 1.5 to 3.5 × 10^8^. These numbers were similar to the number of cells we prepared according to the Mandecki method. With these numbers, DNA gap repair had similar efficiency to the Mandecki method, i.e., 10^−5^-10^−6^ (roughly 1,000 colonies per plating divided by 1.5-3.5 × 10^8^), if all colonies were mutant. However, efficiencies could vary with ssODNs of different homology arm length. In the Mandecki report, the number of mutant colonies using a 15-nucleotide homology arms ssODN was about 10 times higher than a 10-nucleotide homology arms ssODN, and there was only an about 2-fold increase when homology arm length increased from 15 to 30 nucleotides. In DNA gap repair, there was an about 3.5-fold increase when homology arms increased from 10 to 15 nucleotides and about 7-fold increase when homology arms increased from 15 to 30 nucleotides. Thus, significant homology arm size limitation for the Mandecki method occurred between 10 and 15 nucleotides whereas for DNA gap repair this occurred between 15 and 30 nucleotides.

### Comparing background cell contamination in the Mandecki method and DNA gap repair

Linear double stranded DNA cannot transform *E. coli* cells due to lack of non-homologous end joining (NHEJ) pathways and inability to replicate^11, 12^. Due to incomplete digestion, the use of endonucleases to introduce double stranded DNA breaks always leads to some DNA fraction retaining transformation potential and thus contamination of mutants with background colonies. There is also some dependence of the number of background colonies on the method of DNA extraction. Plasmids extracted with non-ionic detergents have lower background colonies than alkali probably because they are free of irreversibly denatured DNA^5, 13^, which is resistant to endonucleases but active in cell transformation. These background colonies are usually present at a frequency of about 10^−4^-10^−5^ in high efficiency electrocomptent cells (10^9^-10^10^ transformants per 1 μg pUC plasmid DNA) at plasmid DNA saturating concentrations. Thus, with DNA gap repair efficiency of 10^−6^, only 1 to 10% of colonies would be expected to be mutant if endonucleases were used for the introduction of double stranded DNA breaks.

PCR products treated with DpnI to remove *in vivo*-derived plasmid DNA template are linear and, as theoretically expected, can establish colonies only when the plasmid is re-circularized as a result of recombination using short homology arms^14^. In Mandecki’s method, restriction endonucleases are used for plasmid linearization, leading to significant background. As shown in Figure 5 of Mandecki’s report, the number of background colonies was about 800. We were unable to reproduce Mandecki’s results with our plasmids and ssODNs, using both 50 and 100 mM CaCl_2_ transformation of JM83 cells. Efficiency of transformation was only 5-10 × 10^4^ transformants per 1 μg pUC plasmid DNA. We also failed to reproduce Mandecki’s results with 10^6^ per 1 μg pUC plasmid DNA JM83 competent cells prepared by Mix&Go protocol and Mix&Go competent cells JM108 (Zymo Research). For all of these (chemically) competent cells there were no colonies with or without a 30-nucleotide homology arm ssODN. In these experiments, the average number of cells treated with DNA was similar to our electroporation experiments, about 3-5 × 10^8^. Thus, for the Mandecki method the background cell frequency can be 10^−5^-10^−6^. In Fig. 4, the frequency of background cells can be determined directly from the colony number formed with the electroporation of targeted DNA, about 10^−7^. Overall, the main feature of Mandecki’s method is that the fraction of mutant colonies is highly variable and depends on experimental conditions (type and location of mutation, the size of targeting ssODNs, and denatured or native targeted DNA).

With our DNA gap repair approach, the wild type sequence is physically removed and PCR preps can be used as targeted DNA for the introduction of different mutations. Thus, very rare intramolecular recombinants of the targeted DNA are the only possible background that can be found both with and without ssODN electroporation^14^. To check this, we sequenced DNA from colonies obtained after co-electroporating native targeted DNA and targeting ssODN into cells with repressed recombineering genes (no heat shock) (Fig. 5). Colony numbers in background and 2×10+1 experiments were similar, and as expected only 14% of colonies were mutant. However, the fraction of mutant colonies increased rapidly for 15-nucleotide homology arms ssODNs, reaching 71%, and becoming 100% for 30-nucleotide homology arms ssODNs (Fig. 5). Out of 11 non-mutant colonies sequenced, all structures were unique except structure 3 (Table 3) which was present in two independent clones. Non-mutant structures were not direct gap end-joining products (structure 0; Table 3). In fact, all of them had gap extension at either the 5’ (structures 3, 5, 8) or 3’ (structures 1-7) gap ends or both (structures 3, 5). Thus, these “non-mutants” were actually mutants with unexpected structures preferentially originating from recombination of one intact end with an internal site located in the other end of the targeting DNA. Among them, there were some produced by recombination of targeted DNA ends with obvious homology (structures 2-5). Others however had no obvious homology between ends (structures 1, 6-8). Interestingly, most of the mutants retained intact 5’-ends terminating in GG (structures 1, 2, 4, 6, 7), and three of them contained deletion of the (GA)4 dinucleotide repeat at the 3’-end (structures 3, 6, 7).

**Table 3.**
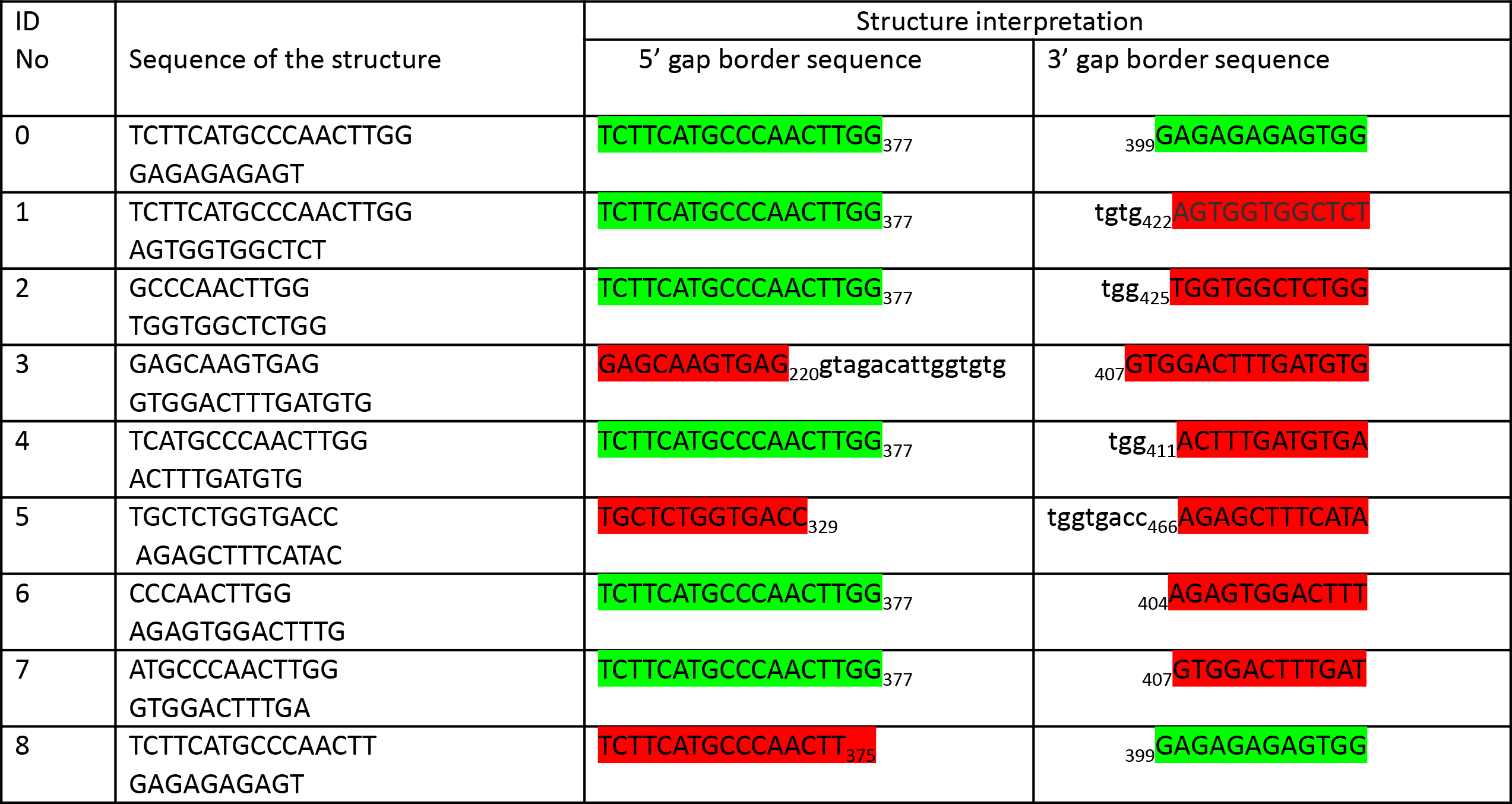
DNA structure of some unexpected mutants from the experiment with native construct DNA/recombineering- and 2×10+1 targeting ssODN reported in Fig. 4. Structure 0 with correct gap boundary sequences are highlighted in green throughout the table. Incorrect gap boundary sequences are highlighted in red. Subscript numbers represent the position of a terminal nucleotide in the wild type sequence (1-714 nucleotides). Sequences in lower case are homological to the other end, presumably causing targeting DNA rearrangements and their deletion from the corresponding structure (second column: sequence of the structure).

To determine the size of terminal direct repeats (TDR) that would generate background cells with a frequency comparable to the efficiency of DNA gap repair, we prepared PCR-derived targeted DNA with TDR of 15 and 24 bp as described in Materials and Methods. With 15 bp terminal repeat, there was more than a 20-fold increase in colony numbers compared to the same DNA without TDR, leading to a frequency of background cells of about 2×10^−5^ (1908/10^8^). There was an additional 30-fold increase in background colony numbers when the size of TDR increased to 24 bp, resulting in a frequency of background cells of about 6×10^−4^ (59,000/10^8^) (Fig. 6). Thus, the frequency of background colonies becomes comparable to the efficiency of DNA gap repair when the size of TDR is in the range of 15-25 bp.

These findings suggest that homology, repeats, and probably GC-rich stretches should be avoided in the ends of the gap to reduce the rate of unexpected mutants. As shown in Fig. 5, with this precaution and increasing the homology arm length of the targeting ssODN, virtually 100% of mutant clones can be produced without contamination of mutants with any other structure. Unlike the Mandecki method, the fraction of both mutants and unexpected mutants are the same for all possible mutations introduced through DNA gap repair because they depend on homology arms and the structure of the ends of targeted DNA.

### Additional Considerations

With the introduction of site specific designer nucleases (ZFN, TALEN, and especially Cas9), DNA repair applications have become standard in DNA engineering partly due to high efficiency, avoiding the need for mutant isolation procedures. Although CRISPR/Cas9 has been successfully used in *E. coli*^15^, its potential application for site-directed mutagenesis appears complex for such a simple model system compared with the relatively straightforward DNA integration approach coupled with selection, our recently described markerless method for the isolation of rare mutants^4, 6^, or our newly described DNA gap repair approach.

## Conclusion

Here we demonstrate that DNA gap repair is independent on lambda red gene product; it does not require targeted DNA denaturation; it functions with targeting DNA homology arms of at least 10 nucleotides and it is potentially background-free. These features are different from recombineering and the Mandecki approach (Table 1), thereby suggesting that these three genetic engineering pathways have at least partly different mechanisms.

## Acknowledgements

DNA sequencing was performed at the University of Nebraska Medical Center DNA Sequencing Core Facility.

## References

1. Copeland, N.G., Jenkins, N.A. & Court, D.L. Recombineering: a powerful new tool for mouse functional genomics. Nature reviews. Genetics 2, 769–779 (2001).

2. Thomason, L.C., Costantino, N., Shaw, D.V. & Court, D.L. Multicopy plasmid modification with phage lambda Red recombineering. Plasmid 58, 148–158 (2007).

3. Mandecki, W. Oligonucleotide-directed double-strand break repair in plasmids of Escherichia coli: a method for site-specific mutagenesis. Proc. Natl. Acad. Sci. U. S. A. 83, 7177–7181 (1986).

4. Lyozin, G.T. et al. Isolation of rare recombinants without using selectable markers for one-step seamless BAC mutagenesis. Nat Methods 11, 966–970 (2014).

5. Lezin, G., Kosaka, Y., Yost, H.J., Kuehn, M.R. & Brunelli, L. A one-step miniprep for the isolation of plasmid DNA and lambda phage particles. PLoS One 6, e23457 (2011).

6. Lyozin, G.T., Kosaka, Y., Bhattacharje, G., Yost, H.J. & Brunelli, L. Direct Isolation of Seamless Mutant Bacterial Artificial Chromosomes. Curr. Protoc. Mol. Biol. 118, 8 6 1–8 6 29 (2017).

7. Court, D.L. et al. Mini-lambda: a tractable system for chromosome and BAC engineering. Gene 315, 63–69 (2003).

8. Yu, D. et al. An efficient recombination system for chromosome engineering in Escherichia coli. Proc. Natl. Acad. Sci. U. S. A. 97, 5978–5983 (2000).

9. Erler, A. et al. Conformational adaptability of Redbeta during DNA annealing and implications for its structural relationship with Rad52. J. Mol. Biol. 391, 586–598 (2009).

10. Hanahan, D., Jessee, J. & Bloom, F.R. Plasmid transformation of Escherichia coli and other bacteria. Methods Enzymol. 204, 63–113 (1991).

11. Bowater, R. & Doherty, A.J. Making ends meet: repairing breaks in bacterial DNA by non-homologous end-joining. PLoS genetics 2, e8 (2006).

12. Rybchin, V.N. & Svarchevsky, A.N. The plasmid prophage N15: a linear DNA with covalently closed ends. Mol. Microbiol. 33, 895–903 (1999).

13. Lezin, G., Kuehn, M.R. & Brunelli, L. Hofmeister series salts enhance purification of plasmid DNA by non-ionic detergents. Biotechnol. Bioeng. 108, 1872–1882 (2011).

14. Zhang, Y., Muyrers, J.P., Testa, G. & Stewart, A.F. DNA cloning by homologous recombination in Escherichia coli. Nat. Biotechnol. 18, 1314–1317 (2000).

15. Jiang, W., Bikard, D., Cox, D., Zhang, F. & Marraffini, L.A. RNA-guided editing of bacterial genomes using CRISPR-Cas systems. Nat. Biotechnol. 31, 233–239 (2013).

